# Epigenetic genes are differentially methylated in the blood of persons with mild cognitive impairment and Alzheimer’s disease

**DOI:** 10.1101/2025.06.18.660344

**Authors:** Samuel G. Faasen, Ligia A. Papale, Alexander Boruch, Phillip E. Bergmann, Kirk J. Hogan, Reid S. Alisch

## Abstract

**Background:** Environmental factors play a role in AD pathology and are mediated by changes in DNA methylation levels.

**Methods and Results:** We investigated whole genome methylation sequencing (WGMS) data from the blood of participants with mild cognitive impairment (MCI, *N*=99), late onset dementia due to Alzheimer’s disease (AD, *N*=109), and who are cognitively unimpaired (CU, *N*=174) to test for differential methylation in 812 genes with roles in epigenetic regulation (*e.g.*, DNA methylation and demethylation, chromatin remodeling, histone modification, and RNA modification) curated from the EpiFactors 2.1 database. 71/812 genes were differentially methylated comprising 190 unique differentially methylated positions (DMPs) in MCI (MCI *vs*. CU pairwise comparison). 60/812 genes were differentially methylated comprising 220 DMPs in AD (AD *vs*. CU pairwise comparison). The majority of differentially methylated genes in both MCI (41/71) and AD (33/60) were histone modification genes and 23 differentially methylated genes were shared in both pairwise comparisons. 96 genes were differentially methylated comprising 243 DMPs between persons with MCI and AD (AD *vs*. MCI pairwise comparison). 10 differentially methylated genes were shared between the 3 pairwise comparisons, including CUGBP elav-like family member 2 (*CELF2*), histone deacetylase 9 (*HDAC9*), RNA binding fox-1 homolog 1 (*RBFOX1*), TATA-box binding protein associated factor 4 (*TAF4*), and thymine DNA glycosylase (*TDG*).

**Conclusion:** Genes that participate in the epigenetic regulation of gene expression, particularly histone modifications, are differentially methylated in blood between persons with and without MCI and AD, warranting further elucidation of their role in the molecular pathogenesis of cognitive decline.

## Background

Late-onset dementia due to Alzheimer’s disease (AD) is characterized by signs and symptoms of dementia that present after the age of 65. Mild cognitive impairment (MCI) is a prodromal stage of AD characterized by mild symptoms with retained functional independence. Eighty percent of persons with MCI will convert to AD within 6 years of their MCI diagnosis (1–3). Evidence indicates that environmental factors increase risk of AD onset and progression through changes in DNA methylation levels [4,5]. DNA methylation is the covalent addition of a methyl group to a cytosine, primarily in a cytosine-phosphate-guanine dinucleotide (CpG), that contributes to regulation of gene expression. Differential DNA methylation in postmortem human brain tissues are reported in genes associated with AD [6,7], with shared patterns of DNA methylation levels between brain and blood [8–11]. Epigenetic mechanisms, including DNA methylation and demethylation, histone modification, and chromatin remodeling, influence AD onset, progression and severity [12–14]. For example, histone modifications are associated with learning, memory, cognition, and synaptic plasticity changes in AD [12,13,15,16]. Further, proteins that regulate chromatin structure and remodeling have altered expression and protein levels in the brains of persons with AD [16,17].

We recently reported differential DNA methylation in blood samples [18,19] from participants with MCI, AD and those who are cognitively unimpaired (CU) from the Wisconsin Alzheimer’s Disease Research Center (WADRC) [20,21] and the Wisconsin Registry for Alzheimer’s Prevention (WRAP) [22]. In the present investigation, we investigated the differentially methylated genes from that data specifically for differentially methylated genes that participate in epigenetic regulation curated from the following 9 categories of the human EpiFactors 2.1 database [23,24]: 1) chromatin remodeling; 2) DNA methylation/demethylation; 3) Histone modifications; 4) Polycomb group [PcG] proteins; 5) protein modification; 6) RNA modification; 7) scaffold proteins; 8) histone subunits; and 9) protein complexes (made up of 234 genes participating in various forms of epigenetic modification).

## Results

### Differential methylation in genes that participate in epigenetic regulation in MCI and AD

#### Differential methylation in MCI

190 DMPs were found in genes that participate in epigenetic pathways between CU and MCI participants (MCI *vs.* CU pairwise comparison; Supplementary Dataset 1). The majority of DMPs were hypermethylated (56%; greater levels of DNA methylation in persons with MCI than in those CU; Fig. 1A). These DMPs annotated to 71 differentially methylated genes and predominantly resided in intronic regions of genes (54%; Fig. 1B) and in genes encoding histone and RNA modifications (46% and 23%, respectively; local false discovery rate [LFDR] < 0.01 and at least one DMP; Fig. 1C). 44/71 genes are involved in histone modification (Fig. 1C). Three differentially methylated genes were histone subunits and 14 were in epigenetic modification complexes, predominantly involved in histone modification (59%; Supplementary Dataset 2).

**Figure 1:**
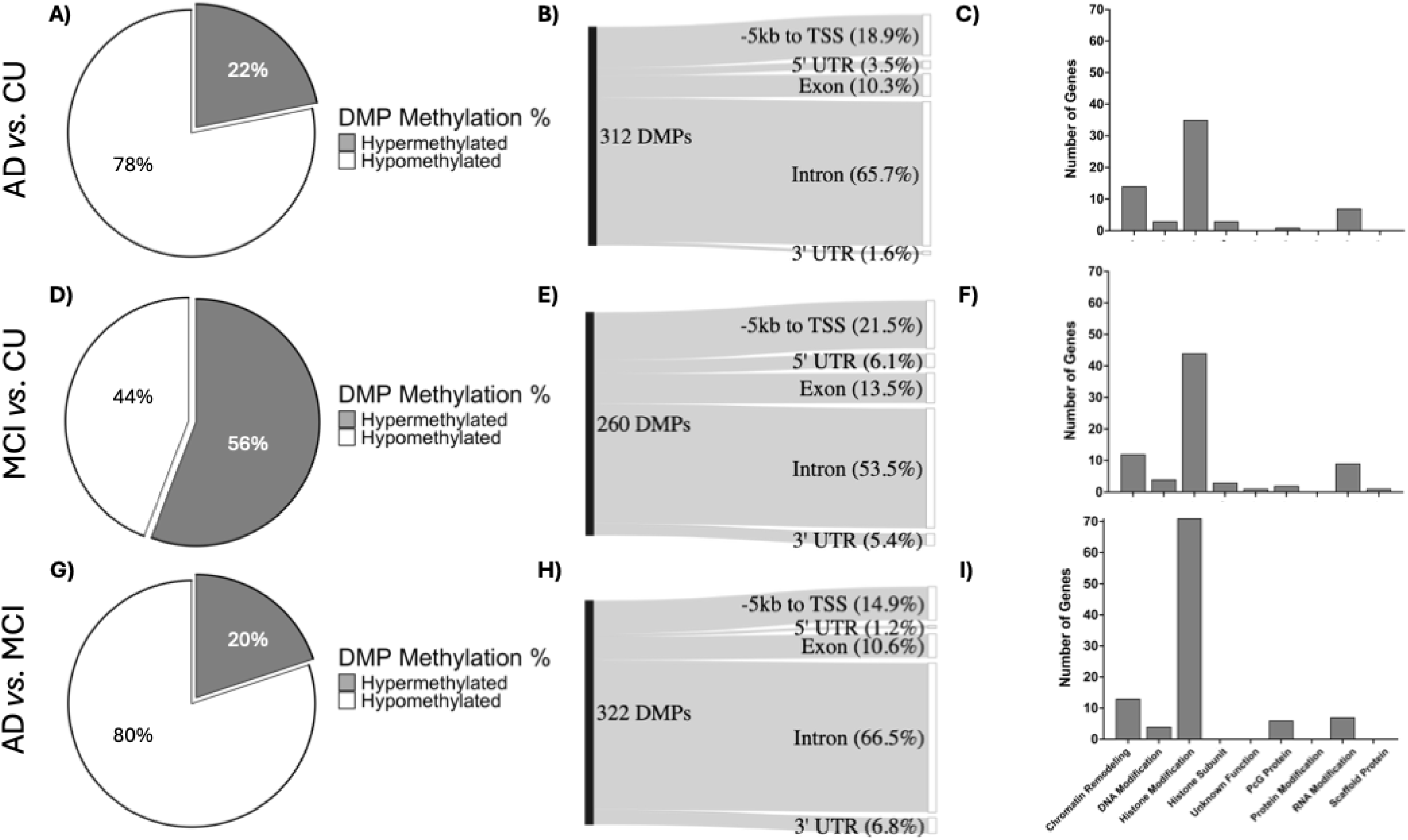
DMPs in genes that encode proteins that participate in epigenetic modification. Pie charts display the proportions of hypermethylated and hypomethylated DMPs in each comparison: MCI *vs*. CU [A]; AD *vs*. CU [D]; and AD *vs*. MCI [G]. Sankey plots show the locations of DMPs in each pairwise comparison relative to genomic structures comprising the following: 5 kilobases (kb) upstream of the transcription start site (TSS; −5kb to TSS); the 5’ untranslated region (5’ UTR); within a gene exon (exon); within a gene intron (intron); and the 3’ untranslated region (3’ UTR): MCI *vs*. CU [B]; AD *vs*. CU [E]; and AD *vs*. MCI [H]. Bar plots show the number of genes from each pairwise comparison, stratified by their broad epigenetic modification types: MCI *vs*. CU [C]; AD *vs*. CU [F]; and AD *vs*. MCI [I].

#### Differential methylation in AD

173 DMPs were in genes that participate in epigenetic pathways between CU and AD participants (AD *vs.* CU pairwise comparison; Supplementary Dataset 1). The majority of DMPs were hypomethylated (78%; greater levels of DNA methylation in persons with CU than in those with AD; Fig. 1D). These DMPs annotated to 60 differentially methylated genes and predominantly resided in intronic regions of genes (66%; Fig. 1E) and in genes encoding histone modifications (56%; local false discovery rate [LFDR] < 0.01 and at least one DMP; Fig. 1F). 35/60 genes are involved in histone modification (Fig. 1C). Three differentially methylated genes were histone subunits, and 14 were in epigenetic modification complexes, mostly involved in histone modification (58%) or chromatin remodeling (39%; Supplementary Dataset 2).

#### Differential methylation between MCI and AD

243 DMPs were in genes that participate in epigenetic pathways between MCI and AD participants (AD *vs.* MCI pairwise comparison; Supplementary Dataset 1). The majority of DMPs were hypomethylated (80%; greater levels of DNA methylation in persons with MCI than in those with AD; Fig. 1G). These DMPs annotated to 96 differentially methylated genes and predominantly resided in intronic regions of genes (66%; Fig. 1H) and in genes encoding histone modifications (61%; local false discovery rate [LFDR] < 0.01 and at least one DMP; Fig. 1I). 71/96 genes are involved in histone modification (Fig. 1C). Zero differentially methylated genes encoded for histone subunits and 30 were in epigenetic modification complexes, mostly involved in histone modification (60%) or chromatin remodeling (27%; Supplementary Dataset 2).

### Shared differential methylation in MCI and AD on genes that participate in epigenetic regulation

#### Shared differentially methylated positions in MCI and AD

23 DMPs were shared between both MCI (MCI *vs.* CU) and AD (AD *vs.* CU); all presented with the same methylation polarity in both comparisons. 19/23 DMPs have decreased methylation levels in both MCI and AD (82%; Fig. 2A). 11 of the shared DMPs were within a single gene, thymine DNA glycosylase (*TDG*). Four of these 11 DMPs were also identified between persons with MCI and AD (AD *vs.* MCI pairwise comparison). All four were located in an exon or the 5’ untranslated region (UTR) of *TDG* (Fig. 2B) and exhibited decreased DNA methylation levels in MCI and AD compared to CU, and increased DNA methylation levels between MCI and AD (Fig. 2C).

**Figure 2:**
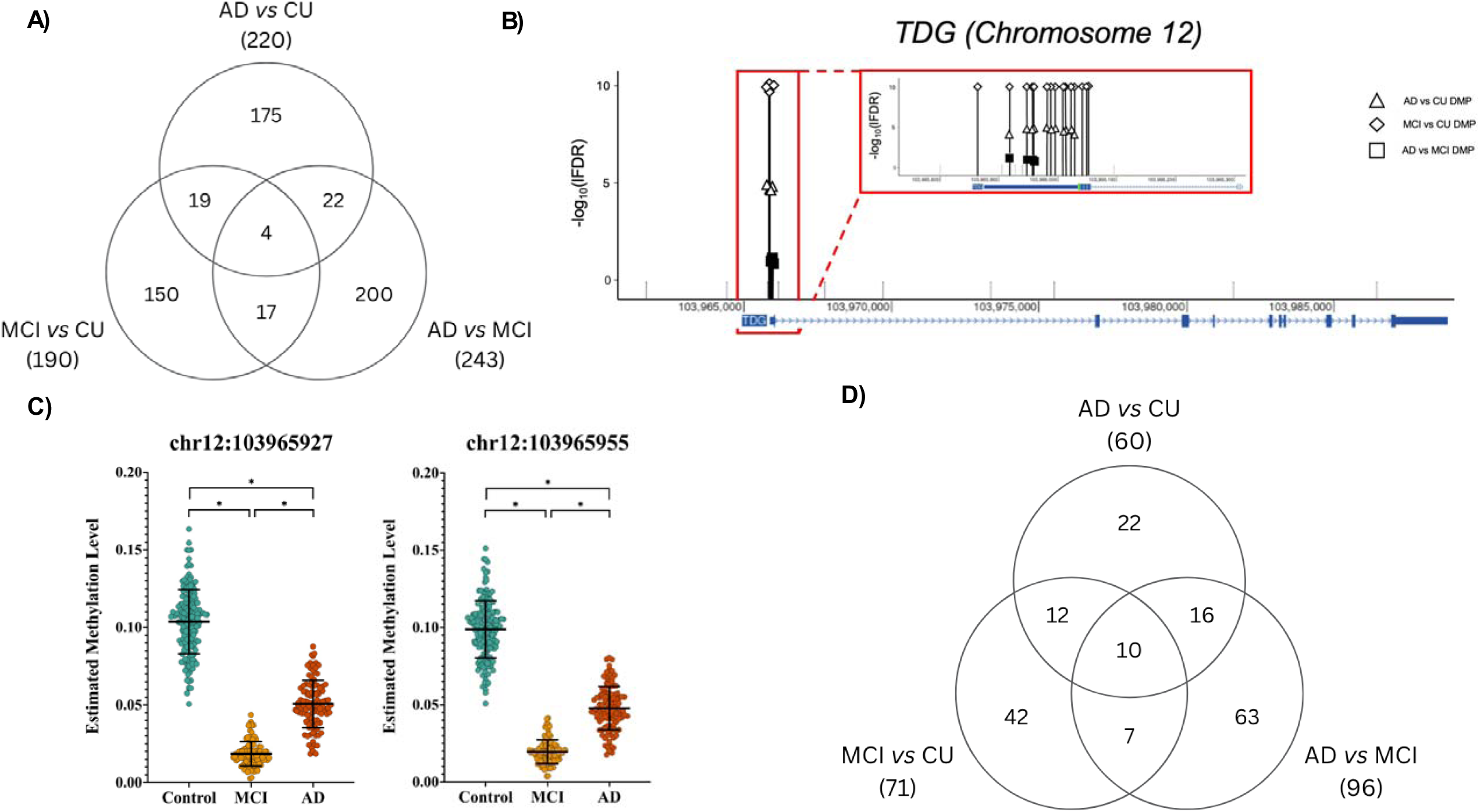
Shared DMPs and genes between the 3 pairwise comparisons. 3-way Venn diagrams showing the overlap of DMPs [A] between the MCI *vs.* CU, AD *vs.* CU, and AD *vs.* MCI pairwise comparisons. Four DMPs within *TDG* are shared between the 3 pairwise comparisons. A gene schematic of the sense strand of *TDG* is depicted [B]. Alignment to human reference genome hg38 coordinates are shown in basepairs (bp) the below the x-axis. Blue rectangles indicate coding exons, and thin blue lines indicate introns. One in 25 non-significant CpGs are shown (grey vertical line). The significance (y-axis) of hypermethylated DMPs (black fill) or hypomethylated DMPs (white fill) are displayed for MCI *vs.* CU (diamond), AD *vs.* CU (triangle), or AD *vs.* MCI (square) pairwise comparisons. A corrected significance level of LFDR < 0.05 and greater than 2.5% methylation difference was adopted for all comparisons. A zoomed-in section of the 5’ UTR of *TDG* is inset into the figure to help visualize the position of the DMPs. Example distributions of the estimated DNA methylation values for 2 of the 4 shared DMPs [C]. Estimated DNA methylation values at a CpG dinucleotide are adjusted for estimated white blood cell proportions, the first two principal components, age, sex, and body mass index (BMI). 3-way Venn diagrams showing the overlap of differentially methylated genes [D] between the MCI *vs.* CU, AD *vs.* CU, and AD *vs.* MCI pairwise comparisons.

#### Differentially methylated genes shared in MCI and AD

23 differentially methylated genes were found in both MCI and AD (Fig. 2D). 11 differentially methylated genes were shared between all three pairwise comparisons, including CUGBP elav-like family member 2 (*CELF2*), histone deacetylase 9 (*HDAC9*), RNA binding fox-1 homolog 1 (*RBFOX1*), TATA-box binding protein associated factor 4 (*TAF4*), and *TDG* (Fig. 2D).

### Differential DNA methylation in enhancer regions interacting with promoters of genes that participate in epigenetic regulation in MCI and AD

#### Differential methylation in enhancers in MCI

41 unique enhancer regions containing 62 DMPs interact with the promoters of 42 genes that participate in epigenetic regulation in MCI (MCI *vs.* CU pairwise comparison; Supplementary Table 3). The majority of DMPs residing in enhancers were hypermethylated (58%; greater levels of DNA methylation in persons with MCI than in those who are CU). 36/42 genes in the EpiFactors database that have promoter-enhancer interactions comprising one or more DMPs have no differential methylation in their gene body (5 kb upstream of TSS to TTS; Supplementary Table 3). Most of these 42 genes encode histone modifications (48%) and chromatin remodeling (34%; Supplementary Table 3).

#### Differential methylation in enhancers in AD

40 unique enhancer regions containing 52 DMPs interact with the promoters of 42 genes that participate in epigenetic regulation in AD (AD *vs.* CU pairwise comparison; Supplementary Table 3). The majority of DMPs residing in enhancers were hypomethylated (70%; greater levels of DNA methylation in persons who are CU than in those with AD). 33/42 genes in the EpiFactors database that have promoter-enhancer interactions comprising one or more DMPs have no differential methylation in their gene body (5 kb upstream of TSS to TTS; Supplementary Table 3). Most of these 42 genes encode histone modifications (47%) or histone subunits (23%; Supplementary Table 3).

#### Differential methylation in enhancers between MCI and AD

69 unique enhancer regions containing 81 DMPs interact with the promoters of 59 genes that participate in epigenetic regulation between persons with MCI and AD (AD *vs.* MCI pairwise comparison; Supplementary Table 3). The majority of DMPs residing in enhancers were hypomethylated (90%; greater levels of methylation in persons with MCI than AD). 41/59 genes in the EpiFactors database that have promoter-enhancer interactions comprising one or more DMPs have no differential methylation in their gene body (5 kb upstream of TSS to TTS; Supplementary Table 3). Most of these 59 genes encode proteins involved in histone modifications (65%; Supplementary Table 3).

#### Shared differential methylation in enhancers in MCI and AD

8 DMPs residing in enhancers were shared between MCI and AD compared to CU, but none were shared between all 3 comparisons (Fig. 3A). 8 genes interacting with enhancer regions containing one or more DMPs were shared between MCI and AD (Fig. 3B). 5 genes interacting with enhancer regions containing one or more DMPs were shared between the 3 comparisons: Bromodomain containing 7 (*BRD7*), Chromodomain helicase DNA binding protein 1 (*CHD1*), IKAROS family zinc finger 1 (*IKZF1*), NIMA related kinase 9 (*NEK9*), and TATA-box binding protein associated factor 9 (*TAF9;* Fig. 3C-3F, Supplementary Fig. 1).

**Figure 3:**
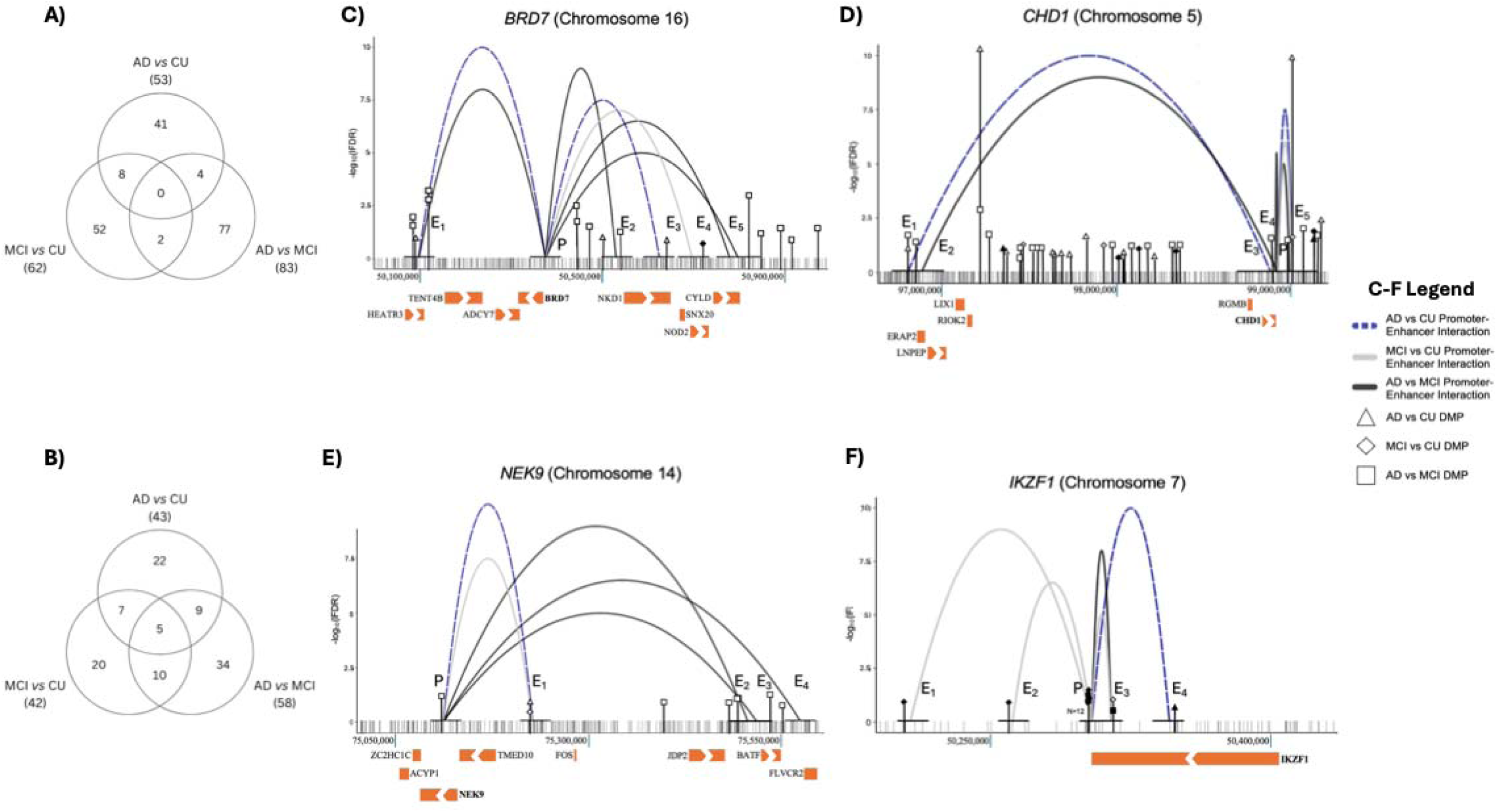
Enhancers comprising unique and shared DMPs between the 3 pairwise comparisons. Three-way Venn diagrams showing the number of unique and shared DMPs within differentially methylated enhancers [A] and unique and shared genes interacting with differentially methylated enhancers [B]. Gene schematic diagrams of 4 genes with interactions with differentially methylated enhancers shared in the 3 pairwise comparisons are shown with gene promoter interactions (arched loops): *BRD7* [C], *CHD1* [D], *NEK9* [E], and *IKZF1* [F]. Alignment to human reference genome hg38 coordinates are displayed in basepairs (bp) below the x-axis. Orange rectangles represent gene bodies, with white arrows inset if the gene body is large enough, to indicate the direction of transcription. The location of the gene name corresponds to the location of the promoter for each gene. Significance (y-axis) of hypermethylated DMPs (black fill) or hypomethylated DMPs (white fill) are displayed for MCI *vs.* CU (diamond), AD *vs.* CU (triangle), or AD *vs.* MCI (square) pairwise comparisons. One in 25 non-significant CpGs are shown for comparison (grey vertical line). Enhancer (E) and promoter (P) regions are designated by black horizontal lines above the grey vertical lines indicating relative positions. Enhancer-promoter interactions are depicted by arched loops for the MCI *vs.* CU (solid light grey), AD *vs.* CU (dashed blue), and the AD *vs.* MCI (solid black) pairwise comparisons. The height of the arched loops is for clarity and does not indicate the significance of each enhancer-promoter interaction. A corrected significance level of LFDR < 0.05 and methylation difference greater than 2.5% was adopted for all comparisons.

## Discussion

We investigated published WGMS data that provided a panel of differentially methylated positions and genes associated with MCI and AD (18,19] for differential methylation in 812 genes that participate in epigenetic regulation curated from the EpiFactor database. 56% of DMPs in genes involved in epigenetic modification were more methylated in MCI than in CU participants; in contrast, 22% of DMPs in genes were more methylated in AD than in CU participants. The majority of DMPs found between MCI and AD participants in genes involved in epigenetic modifications also had lower DNA methylation levels in AD, suggesting a trend of increasing DNA methylation levels from CU to MCI, and decreasing DNA methylation levels from MCI to AD in genes involved in epigenetic modifications. Since gains of DNA methylation are linked to decreased gene expression [25], these findings suggest the Epifactors database comprises genes with broad transcriptional repression in MCI participants compared to those with AD.

Histone modifications comprise the most differentially methylated genes in each of the 3 pairwise comparisons. Changes in the acetylation and/or methylation at diverse lysine and threonine histone residues have been previously reported in blood and postmortem AD brain tissues [12,13,15,26]. 8/10 histone modification writers differentially methylated in AD (AD *vs.* CU) have amino acid targets that also show altered acetylation, methylation, or phosphorylation levels in postmortem brain tissue from persons with AD [12]. Histone modification complexes contained the most differentially methylated genes of all types of complexes in the 3 pairwise comparisons. Validation of these results is necessary.

Ten genes are shared between all 3 pairwise comparisons, including CUGBP elav-like family member 2 (*CELF2*), histone deacetylase 9 (*HDAC9*), RNA binding fox-1 homolog 1 (*RBFOX1*), TATA-box binding protein associated factor 4 (*TAF4*), and Thymine DNA glycosylase (*TDG*). *CELF2*, as a member of the CELF family, plays various roles in co-transcriptional and post-transcriptional RNA processing, stability, and translation. This gene is a GWAS identified risk factor for AD [27] and has a direct role in the metabolism of Tau proteins [28]. *HDAC9* is a histone deacetylase. *HDAC9* function in humans induces memory impairment, increases amyloid-β deposition, promotes tau hyperphosphorylation, increases neuroinflammation and mitochondrial dysfunction in AD and with age [29–31]. *RBFOX1* participates in tissue-specific splicing of mRNA precursors. *RBFOX1* is a key susceptibility gene in the balance between resilient and phenotype-presenting AD populations [32,33] and has loci associated with a higher AD incidence level in older Black adults [34], increased somatic mutations with age [35], and amyloidosis [36]. Reduced expression of *RBFOX1* is correlated to higher amyloid-β burden and worse cognition during life [36]. *TAF4* is a core subunit of the transcription factor II D (TFIID) complex, is required for the formation of the RNA polymerase II pre-initiation complex, and is essential for embryogenesis and embryonic stem cell differentiation [37]. *TDG* controls the base excision repair of guanine/thymine DNA mismatches, created by the deamination of 5-methylcytosine, back to guanine/cytosine base pairs, and is essential to the development and maintenance of chromatin states by recruiting chromatin-modifying enzymes and protecting CpG-rich promoter sequences from hypermethylation [38]. Alterations to *TDG* would have global consequences due to its major role in protecting the stability of the genome and epigenome [39]. Differential methylation in these genes may cause vast alterations to the epigenome and transcriptome.

Certain gene families were not differentially methylated in the present data, including the DNA methyltransferases (DNMT) and ten-eleven translocase (TET) families of DNA methylation and demethylation enzymes, except for *DNMT1* in MCI and AD, *DNMT3B* in AD, and *TET2* in MCI. In turn, most histone deacetylases assessed using the EpiFactors dataset are not differentially methylated in the three pairwise comparisons (AD *vs.* CU, 7/9; MCI *vs.* CU, 7/9; AD *vs.* MCI, 6/9). The lack of differential methylation in these families of genes suggests that the DNA methylation changes we observe are gene and pathway specific.

We identified DMPs in enhancer regions known to interact with the promoters of genes that participate in epigenetic regulation, including genes not found to be differentially methylated in any pairwise comparisons, thereby increasing the aggregate number of genes with altered DNA methylation levels in persons with MCI and AD. The promoters of genes encoding histone modifications represented the most targeted by enhancer regions containing one or more DMPs (47-65%). Five genes shared the interaction with enhancer regions containing one or more DMPs in all three pairwise comparisons: Bromodomain containing 7 (*BRD7*), Chromodomain helicase DNA binding protein 1 (*CHD1*), IKAROS family zinc finger 1 (*IKZF1*), NIMA related kinase 9 (*NEK9*), and TATA-box binding protein associated factor 9 (*TAF9*). *BRD7* acts as a transcriptional coactivator or corepressor as a component of chromatin remodeling complexes [40], is involved in synapse formation and oligodendrocyte differentiation and myelination [41,42] and has been linked to apoptosis in AD [43]. *CHD1* encodes a protein that regulates the opening of chromatin and plays a role in transcription [44] and has been identified as a key transcriptional regulator of switch genes in the brains of AD patients [45]. *IKZF1* encodes a hematopoietic zinc finger transcription factor participating in gene expression via chromatin remodeling [46,47]. *IKZF1* is specifically expressed in adult microglia in the human cortex and hippocampus, and its loss has been found to induce spatial learning deficits, impaired hippocampal long-term potentiation, abnormal microglial morphology, astrogliosis, and composition changes to the inflammasome and synaptosome [48]. *IKZF1* has increased expression in postmortem hippocampal samples from AD patients [48]. *NEK9* is a serine-threonine protein kinase heavily involved in mitosis and has been identified as a potential therapeutic target in AD due to its roles in tauopathy and tau phosphorylation [49]. *TAF9* is one member of the family of TAFs required for activated transcription. Studies have identified *TAF9* as a top novel target for diagnosis and therapy in AD [50]. Together, these data identify DMPs in enhancer regions known to interact with the promoters of key genes that participate in epigenetic regulation in MCI and AD.

Future work must validate these findings, investigate more genes involved in epigenetic modification not currently listed in EpiFactors, and integrate these results with other omics data to improve the molecular resolution of AD pathology.

## Conclusions

Genes that participate in the epigenetic regulation of gene expression are themselves differentially methylated in blood between persons with and without MCI and AD. Differentially methylated positions (DMPs) within these genes stand to further elucidate the molecular pathogenesis of MCI and AD and may enhance the utility and accuracy of contemporary plasma biomarkers.

## Methods

Details of the cohorts, sample acquisition and handling, DNA methylation sequencing and data analysis have been previously published [18,19]. Sections describing both study participants and original differential DNA methylation analysis are summarized below.

### Study participants

This research was conducted in accord with the Declaration of Helsinki. The experimental protocol was approved by the institutional review board (IRB) of the University of Wisconsin School of Medicine and Public Health, Madison, Wisconsin. All participants signed an IRB-approved informed consent. Participants were enrolled in the Wisconsin Alzheimer’s Disease Research Center (WADRC) [20,21], or the Wisconsin Registry for Alzheimer’s Prevention (WRAP) [22]. Participants were classified into three groups based on cognitive status: 109 participants with Alzheimer’s disease (AD), 99 with mild cognitive impairment (MCI), and 174 who are cognitively unimpaired (CU). Participants are evaluated annually for cognitive status with cognitive performance tests determined by a consensus conference panel based on National Institute on Aging–Alzheimer’s Association (NIA-AA) criteria. [51]

### Whole genome methylation sequencing acquisition and analysis

High molecular weight genomic DNA was extracted from blood samples acquired on the nearest appointment date following MCI or AD diagnosis. The DNA samples underwent whole genome methylation sequencing (WGMS), and the resulting data was processed using standard procedures. A CpG was defined as a differentially methylated position (DMP) if it has a local false discovery rate (LFDR) < 0.05 and differential methylation > 2.5% between groups. Genes were considered differentially methylated if they contained at least one DMP and had a gene-wide LFDR < 0.01. Genomic coordinates for all protein coding genes were obtained from ENSEMBL (v86) [52]. Genes were defined as sequences spanning 3 kilobases (kb) 5′ of a transcription start site (TSS) to 200 base pairs (bp) 3′ of a transcription termination site (TTS). This approach generated 9,756 DMPs and 1,743 differentially methylated genes between CU and MCI, 14,530 DMPs and 1,847 differentially methylated genes between CU and AD, and 19,759 DMPs and 2,929 differentially methylated genes between MCI and AD participants in the previous publication [19].

### Selection of Genes Encoding Proteins that Participate in Epigenetic Regulation of Gene Expression: The *EpiFactors 2.1 Database*

EpiFactors is a useful database that specifically catalogs human proteins and complexes with a role in epigenetic regulation, generated from reputable external databases, and provides a wealth of information and data on epigenetic factors, including expression levels across human primary cell samples, cancer cell lines, and post-mortem tissues. This database is useful for researchers studying epigenetic contributions to complex human disease [23,24]. The first dataset contained genes encoding proteins directly participating in epigenetic modifications.

These genes fell into a number of categories: genes with regulatory roles in chromatin remodeling (*i.e.*, chromatin remodeling, chromatin remodeling cofactor), DNA methylation/demethylation (*i.e.*, DNA modification, DNA modification cofactor), Histone modification (*i.e.*, histone chaperone, histone modification, histone modification cofactor, histone modification erase, histone modification erase cofactor, histone modification read, histone modification read cofactor, histone modification write, histone modification write cofactor), Polycomb group (PcG) protein, protein modification, RNA modification, scaffold protein, and transcription factors. Transcription factors were excluded from this paper for exploration in a future study. In total, 788 genes were obtained for downstream analysis from this dataset. The second dataset contained 101 genes encoding histone subunits. The third dataset contained 73 protein complexes, made up of 368 genes, participating in various forms of epigenetic modification. The genes utilized in these complexes may already exist in the first dataset. Combining all three datasets generated a list of 1,017 genes for differential methylation analysis, 812 of these genes were contained in the Madrid *et al.* dataset.

### Identification of Differentially Methylated EpiFactors genes

Gene and DMP files from the previous publication [19] were utilized for EpiFactors analysis, at the same cutoffs (DMP: LFDR < 0.05 and differential methylation > 2.5%; gene: gene-wide LFDR < 0.01 and 1+ DMP). DMPs were annotated to genomic structures with annotatr (version 1.28.0) [53] for comparison overlapping, genic region and methylation polarity analyses. Differentially methylated genes that encode proteins that participate in epigenetic regulation of gene expression (henceforth just called differentially methylated genes) were matched between the *EpiFactors 2.1 Database* master list and differentially methylated genes identified in the previous publication (*i.e.*, MCI *vs.* CU, AD *vs.* CU, AD *vs.* MCI).

### Differential DNA methylation analysis of blood promoter-enhancer interactions

The genomic coordinates obtained from ENSEMBL (v86) [53] were extended to include 3 kilobases 5’ of a TSS and 200 base pairs 3’ of a transcription termination site, and promoters were updated to be defined as regions 5 kb upstream of a TSS and 200 bases downstream of a TSS. Differential DNA methylation levels at promoter-enhancer loci and their interactions in white blood cells were examined using published Promoter Capture Hi-C (PCHi-C) data [54]. An interaction was deemed significant if the median ChiCAGO score across all blood cell types was > 5. Interactions were then filtered, false positives (*e.g*., bait-bait) and sex chromosome interactions were removed, and data lifted over to the GRCh38 human genome reference version with the UCSC liftOver tool [55]. An enhancer was considered differentially methylated if it contained one or more DMPs. This approach yielded 104,554 promoter-enhancer interactions comprising 58,654 distinct enhancers interacting with 9,631 distinct promoters of genes throughout the genome. Differentially methylated enhancers (defined as an enhancer comprising 1+ DMP) were then limited to those interacting with the promoters of genes identified in the EpiFactors master list (above).

### Software for statistical analysis and data visualization

All statistical analyses (*i.e.* DMP and differentially methylated gene generation and identification, EpiFactors overlap, and statistical significance analysis) were conducted in R (v4.2.0). Sankey, Venn diagram, and enhancer-promoter figures were generated with ggplot2 (version 3.4.2) [56], networkD3 (version 0.4) [57], and stats (version 3.6.2) [58]. Pie charts, bar plots and line graphs were constructed using *GraphPad Prism* (version 10.2.3) [59].

## Declarations

### Ethics approval and consent to participate

The experimental protocol was approved by the Institutional Review Board (IRB) of the University of Wisconsin School of Medicine and Public Health, Madison, WI. All participants signed an IRB-approved informed consent.

### Consent for publication

Not applicable.

### Availability of data and materials

The datasets supporting the conclusions of this article are included within the article (and its additional files).

### Authors’ Contributions

LRC acquired and provided the participants’ blood. LAP extracted the DNA from the blood. SK analyzed the data and interpreted the general trends. SGF further analyzed and interpreted all the data presented, as well as wrote the manuscript. KJH, RSA, and LAP all contributed to the conception and design of the project, as well as the writing of the manuscript. All authors read and approved the final manuscript.

## Acknowledgements

The authors would like to acknowledge and thank the participants and study personnel who made this work possible.

## Abbreviations

AD: Alzheimer’s disease
BMI: Body mass index
BRD7: Bromodomain containing 7
CELF2: CUGBP elav-like family member 2
CHD1: Chromodomain helicase DNA binding protein 1
CpG: Cytosine-guanine dinucleotide
CU: Cognitively unimpaired
DMP: Differentially methylated position
DNA: Deoxyribonuleic acid
DNMT1: DNA methyltransferase 1
DNMT3A: DNA methyltransferase 3A
DNMT3B: DNA methyltransferase 3B
EPI: Enhancer-promoter interaction
HDAC9: Histone deacetylase 9
IKZF1: IKAROS family zinc finger 1
LFDR: Local false discovery rate
kb: kilobases
MCI: Mild cognitive impairment
NEK9: NIMA related kinase 9
NIA-AA: National Institute on Aging-Alzheimer’s Association
PcG: Polycomb group
PCHi-C: Promoter Capture Hi-C
RBFOX1: RNA binding fox-1 homolog 1
RNA: Ribonucleic acid
TAF: TATA-box binding 379 protein associated factor
TAF4: TATA-box binding protein associated factor 4
TAF9: TATA-box binding protein associated factor 9
TDG: Thymine DNA glycosylase
TET: Tet methylcytosine dioxygenase
TET2: Tet methylcytosine dioxygenase 2
TSS: Transcription start site
TTS: Transcription termination site
WADRC: Wisconsin Alzheimer’s disease research center
WGMS: Whole genome methylation sequencing
WRAP: Wisconsin registry for Alzheimer’s prevention

