## Supplemental Figure 1 for "Epigenetic genes are differentially methylated in the blood of persons with mild cognitive impairment and Alzheimer’s disease"

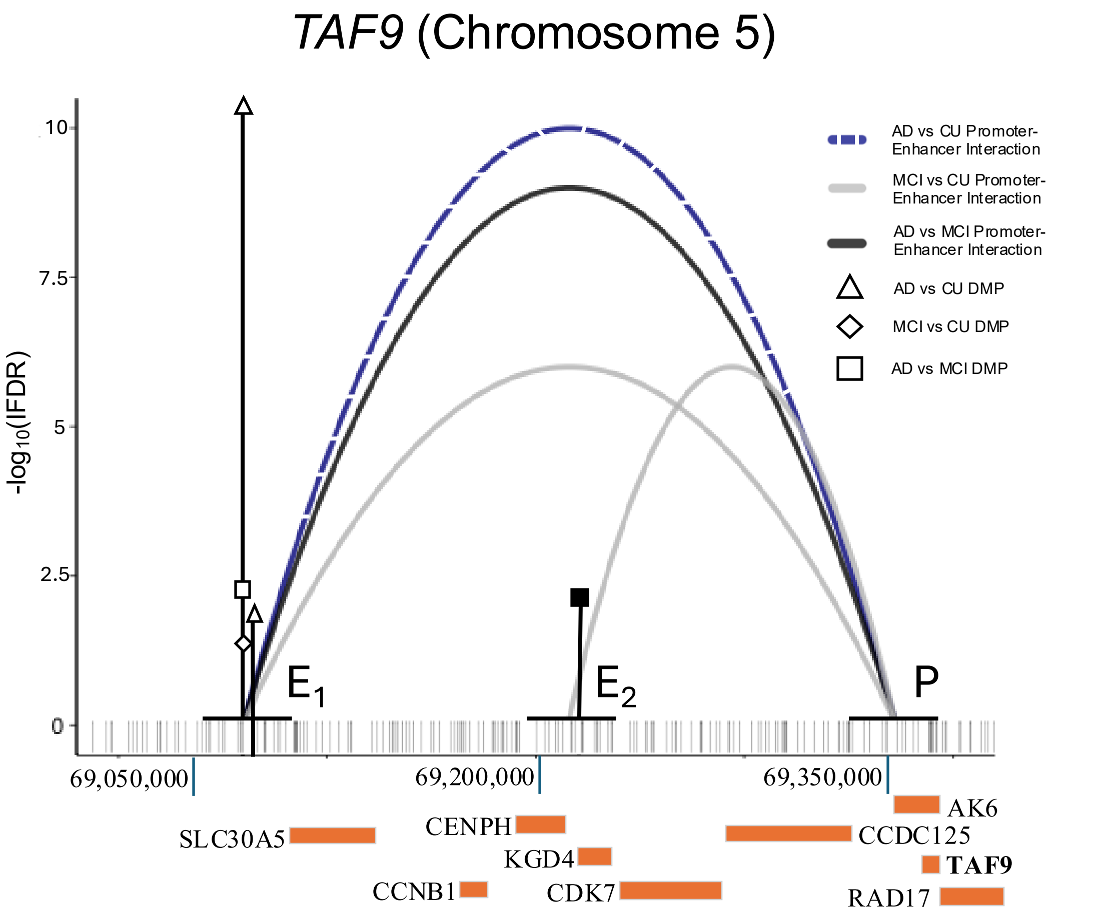


**Supplemental Figure 1:** Enhancers comprising unique and shared DMPs between the 3 pairwise comparisons. Gene schematic diagrams of a gene with interactions with differentially methylated enhancers shared in the 3 pairwise comparisons is shown with gene promoter interactions (arched loops). Alignment to human reference genome hg38 coordinates are displayed in basepairs (bp) below the x-axis. Orange rectangles represent gene bodies, with white arrows inset if the gene body is large enough, to indicate the direction of transcription. The location of the gene name corresponds to the location of the promoter for each gene. Significance (y-axis) of hypermethylated DMPs (black fill) or hypomethylated DMPs (white fill) are displayed for MCI vs. CU (diamond), AD vs. CU (triangle), or AD vs. MCI (square) pairwise comparisons. One in 25 non-significant CpGs are shown for comparison (grey vertical line). Enhancer (E) and promoter (P) regions are designated by black horizontal lines above the grey vertical lines indicating relative positions. Enhancer-promoter interactions are depicted by arched loops for the MCI vs. CU (solid light grey), AD vs. CU (dashed blue), and the AD vs. MCI (solid black) pairwise comparisons. The height of the arched loops is for clarity and does not indicate the significance of each enhancer-promoter interaction. A corrected significance level of LFDR < 0.05 and methylation difference greater than 2.5% was adopted for all comparisons.
